# PhylSim: A New Phylogenetic Tool for Modeling Trait Evolution and Speciation

**DOI:** 10.1101/2020.05.13.094706

**Authors:** P. Spivakovsky, P. Lio, M. S. Clark

## Abstract

We present a new software tool in R, the PhylSim package, which allows the user to simulate trait evolution on a phylogeny, while varying the speciation rate and evolutionary model over the course of the simulation. In particular, the user can specify any number of regimes, with a different speciation rate for each regime, or set the speciation rate to vary when the value of a trait exceeds a certain threshold. The evolutionary model can also be varied as many times as necessary over the course of the simulation, to account for traits tending toward different optima at different periods in evolutionary history. We illustrate the use of the package with an example, a simulation of the adaptive radiation of Antarctic notothenioids in the Southern Ocean.

## 1 Introduction

Understanding trait evolution and speciation over time are important problems in modern biology and bioinformatics. Considerable work has been done in this area by previous authors. For example, the evolution of a numerical trait over a phylogeny has been considered previously by Butler and King (2004) [1]. In this work, the authors model random evolution of traits using an Ornstein-Uhlenbeck diffusion process occuring on a species tree over time. Their implementation resulted in the OUCH package [1].

Bartoszek et al. (2012) [2] then extended this to consider multiple traits evolving together on the species tree. The power of this multivariate approach is that it can consider interactions occurring between multiple traits evolving simultaneously. In particular, a vector of trait values 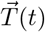 can be subdivided into two trait vectors, 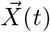 and 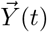, representing “effect” and “response” traits, to model different types of trait interactions; for example, cases of co-adaptation between traits, or some traits responding to the effects of others. This approach was implemented in the mvSLOUCH package, which allowed greater flexibility than the OUCH package for testing evolutionary hypotheses over a phylogeny.

The mvSLOUCH framework was further improved by Bartoszek and Lio (2019) [3] with the ‘pcmabc’ package. This package uses Approximate Bayesian Computation to fit parameters of the stochastic process, thereby making the computation more efficient. It also relies on the ‘yuima’ package [4] for solving SDEs more robustly.

Both the mvSLOUCH and ‘pcmabc’ packages allow the user to simulate trait evolution on a phylogeny under a particular evolutionary model and speciation rate. In the case of the ‘pcmabc’ package, even switching between rates is allowed by the simulation. However, neither package allows the user to specify a large number of regimes, with different evolutionary models and speciation rates for each regime. This additional functionality is provided by the PhylSim package, presented here.

With this extension, PhylSim allows greater flexibility in testing hypothesis of evolution and adaptation in situations where speciation rate is suspected to have varied significantly over geological time. As a result, the package could be useful for studying phylogenetic dynamics under changing environmental conditions, cases of adaptive radiation, and other scenarios.

This paper is organised as follows. In Section 2, we present the PhylSim package and explain the basics of how it works. In Section 3, we use the package to simulate the adaptive radiation of Antarctic notothenioids in the Southern Ocean, as an example. In Section 4, we consider a simulation using multiple trait values. In Section 5, we consider parameter estimation using PhylSim. Finally, in Section 6, we summarize our conclusions and present directions for future work.

## 2 The PhylSim Package

The PhylSim package simulates the evolution of traits on a phylogeny, allowing the user to freely vary speciation rates and evolutionary models over the course of the simulation. PhylSim makes use of the ‘pcmabc’ and ‘yuima’ packages in R, so it can interpret any of the SDE models accepted by ‘yuima’, ie. single and multivariable diffusion processes, Brownian motion, Ornstein-Uhlenbeck with and without jumps, etc.

The user only needs to specify which model to use for each regime, and the times at which the regimes change (arbitrarily many changes are allowed by the package). The regimes for the evolutionary models need not match the regimes for the speciation rates, in order to give the user more flexibility.

At the branch level, PhylSim uses the function ‘simulate sde on branch’ from ‘pcmabc’ package to simulate trait evolution on branch segments. The length of these segments can be specified by the user. In each segment, the trait evolves according to the evolutionary model corresponding to that time period (depending on the regime that segment is in). Also, the probability of branching occurring in a given segment depends on the speciation regime for that segment. Thus, the speciation rate is time dependent, and also trait dependent, as the user can set the rates to change in a given regime when the value of the trait exceeds a certain threshold.

The schematic diagram in Figure 1 summarizes the basics of how PhylSim works:

**Figure 1:**
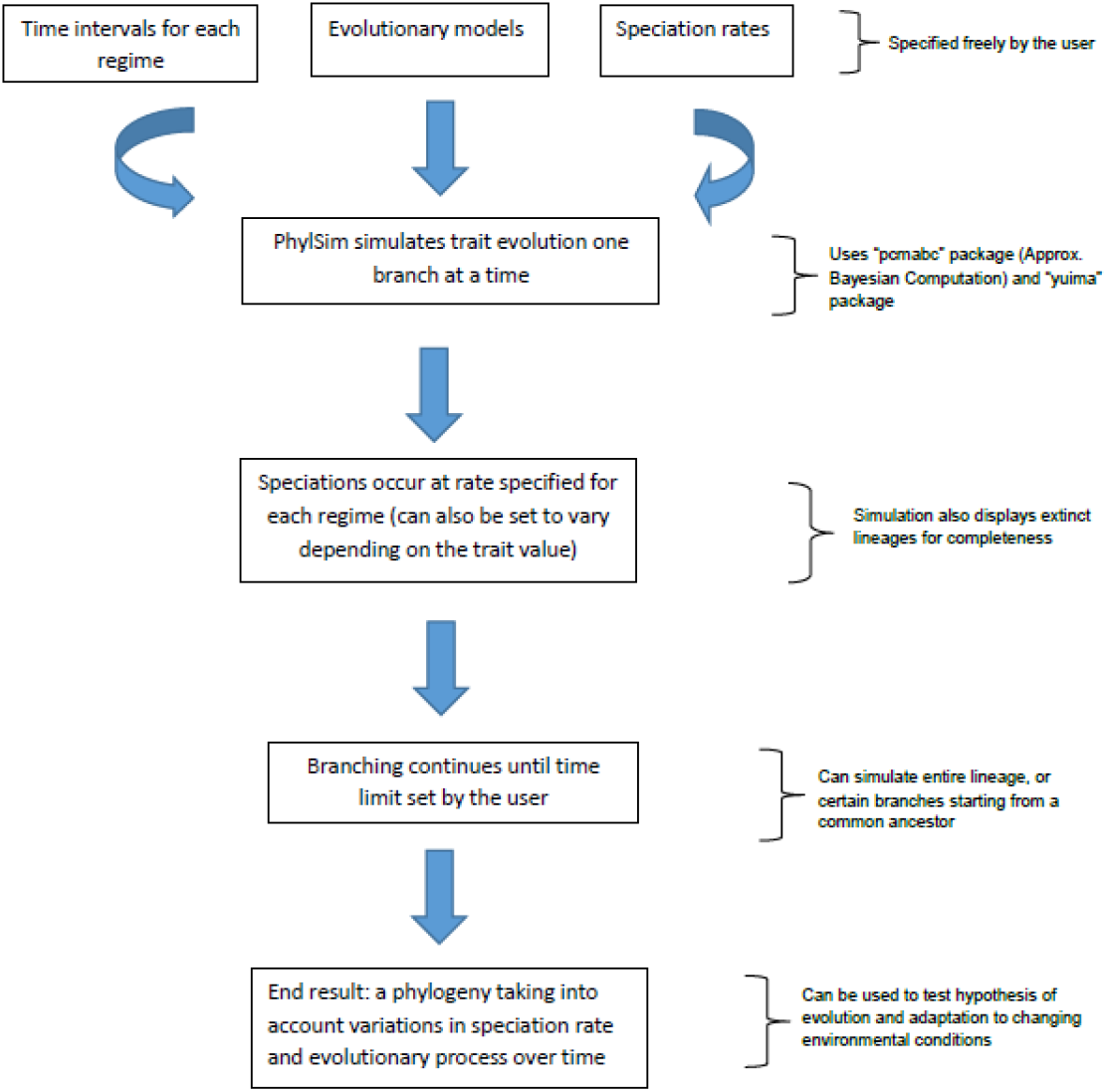
Diagram showing the basics of the PhylSim package.

Also, an example of the early stages of a simulation with an arbitrary numerical trait is shown in Figures 2 through 4. In Figure 4, the speciation rate is set to increase after *t* = 10, which results in increased branching after that time.

**Figure 2:**
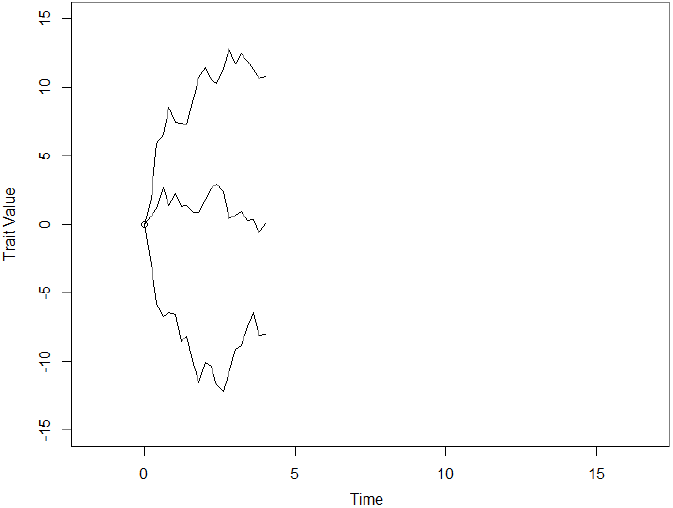
Simulation until *t* = 4, with trait value following an Ornstein-Uhlenbeck process with jumps.

**Figure 3:**
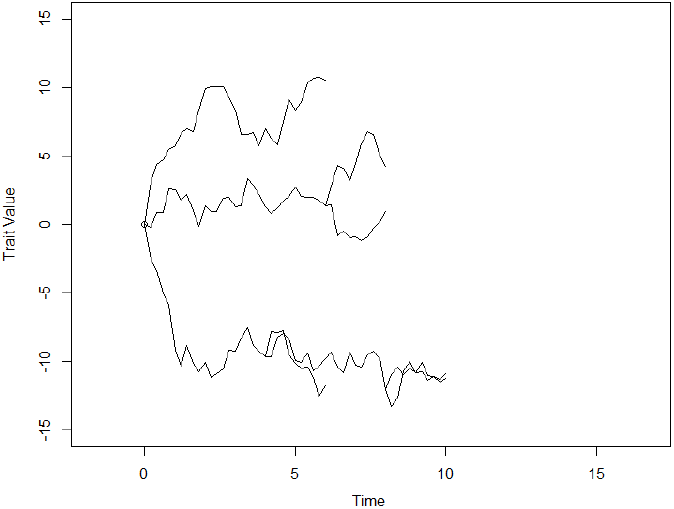
Simulation until *t* = 10, with trait value following an Ornstein-Uhlenbeck process with jumps.

**Figure 4:**
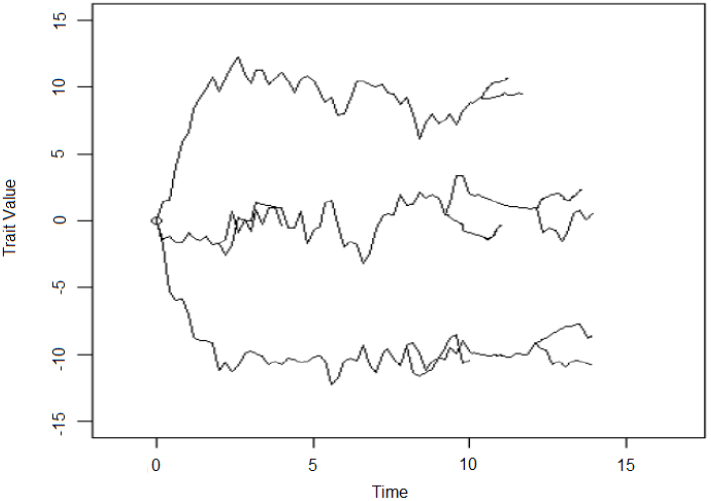
Simulation until *t* = 15, with increased speciation rate after *t* = 10.

The main function in the PhylSim package is called as follows:

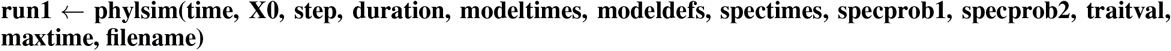

Here, ‘time’ is the starting time of the simulation, and ‘X0’ is the initial value (in the single variable case; it can also be a vector of trait values for the multivariable case). The argument ‘step’ is the step size for SDE solving throughout the simulation.

Next, ‘modeltimes’ is a vector of times indicating when to change the evolutionary model of the simulation (regime changes); this vector can be of any length specified by the user, provided it is of the same length as ‘modeldefs’. The argument ‘modeldefs’ indicates which evolutionary model to use for each regime (the model names are passed as strings). Each model name must be defined previously, in the same way as in the ‘yuima’ package. A typical model definition would be

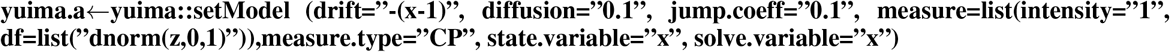

for a model with drift, diffusion and random jumps in the value of the trait. Next, the argument ‘spectimes’ is a vector of times indicating when to change the speciation rate regime. The argument ‘traitval’ is a vector of threshold trait values; if a trait exceeds the threshold value in its regime, the speciation rate is set to the value given by ‘specprob2’, otherwise the default is given by ‘specprob1’.

PhylSim follows Cox’s method from the ‘pcmabc’ package, and relies on the ‘yuima’ package for robust solution of stochastic differential equations on each branch of the phylogeny.

## 3 Simulation of An Adaptive Radiation

As an example, we consider the adaptive radiation of Antarctic notothenioids in the last 35 million years, during the period of cooling of the Southern Ocean. The trait value we consider is the number of copies of a protein kinase gene, Prkg1-201, which is found in all Antarctic notothenioids and their temperate relatives.

Since Prkg1-201 is a key mediator in the nitric oxide cycle, this gene is believed to have been preferentially duplicated in Antarctic notothenioids in reponse to oxidative stress as the climate cooled [7]. The number of copies in each species is the value to which the Ornstein-Uhlenbeck process tends over time.

The Ornstein-Uhlenbeck process is governed by the following stochastic differential equation:

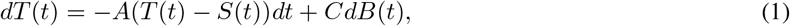

where *T* (*t*) is the value of the trait at time *t*; *A* is the genetic drift term; *B*(*t*) is a standard Brownian motion; *S*(*t*) is the trait distribution to which the process tends; and *C* is a stochastic diffusion term [1]. The solution to the above differential equation is of the form

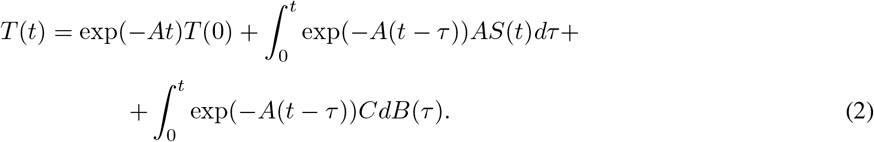

The Antarctic species we consider are: *Parachaenichthys charcoti* (Antarctic dragonfish) [16]; *Dissostichus mawsoni* (Antarctic toothfish) [17]; *Notothenia coriiceps* (Antarctic bullhead notothen) [18]; *Chaenocephalus aceratus* (blackfin icefish) [19]; *Chionodraco myersi* (Myer’s icefish) [20]; *Pseudochaenichthys georgianus* (South Georgia icefish) [23]; and *Harpagifer antarcticus* (Antarctic spiny plunderfish) [23].

The temperate species are: *Eleginops maclovinus* (Patagonian robalo) [17]; *Bovichtus variegatus* (New Zealand thornfish) [21]; *Bovichtus diacanthus* (Tristan klipfish) [22]. These species were chosen based on the availability of genetic sequence data, and to be broadly representative of the main lineages of Antarctic and temperate notothenioids.

We simulate the change in the environmental conditions in this case by setting three different regimes. The first regime, from *t* = 0 to *t* = 12 Myrs, corresponds to a period of relatively stable temperatures in the Southern Ocean around 8 degrees Celsius, as described by Crame (2018) [5]. During this period, the simulation follows an Ornstein-Uhlenbeck process with a moderate speciation rate due to the climate conditions.

The second regime is a period of warming of the Southern Ocean, between *t* = 12 and *t* = 18 Myrs, in which the temperature increases to about 12 degrees [5]. During this time, the speciation rate is significantly reduced, and there are smaller jumps in the value of the trait in the Ornstein-Uhlenbeck process.

The third regime, from *t* = 18 to *t* = 35 Myrs (ie. the present day, since the simulation starts at 35 million years ago), corresponds to the cooling of the Southern Ocean to current temperatures, with increased glacial cycles in the last 5 million years [5]. During this time, the speciation rate increases significantly, not only as a result of the change in temperature, but also due to repeated fragmentation of breeding populations caused by advance and retreat of the glaciers (the so-called biodiversity pump) [7].

The results of the simulation are shown in Figure 5. We observe that the increased rate of speciation in the last 5-10 million years is clearly reflected in the simulation, with a notable increase in the rate of branching during that regime. Note that extinct lineages are also shown in the simulation for completeness. Antarctic species are labeled in black, related temperate species are labeled in red. Time is in millions of years. These results agree with the findings of Near at al. (2012) [8], which suggest that although antifreeze proteins evolved in Antarctic fish more than 20 million years ago, the main diversification of species occurred only in the last 5-10 million years of their evolution.

**Figure 5:**
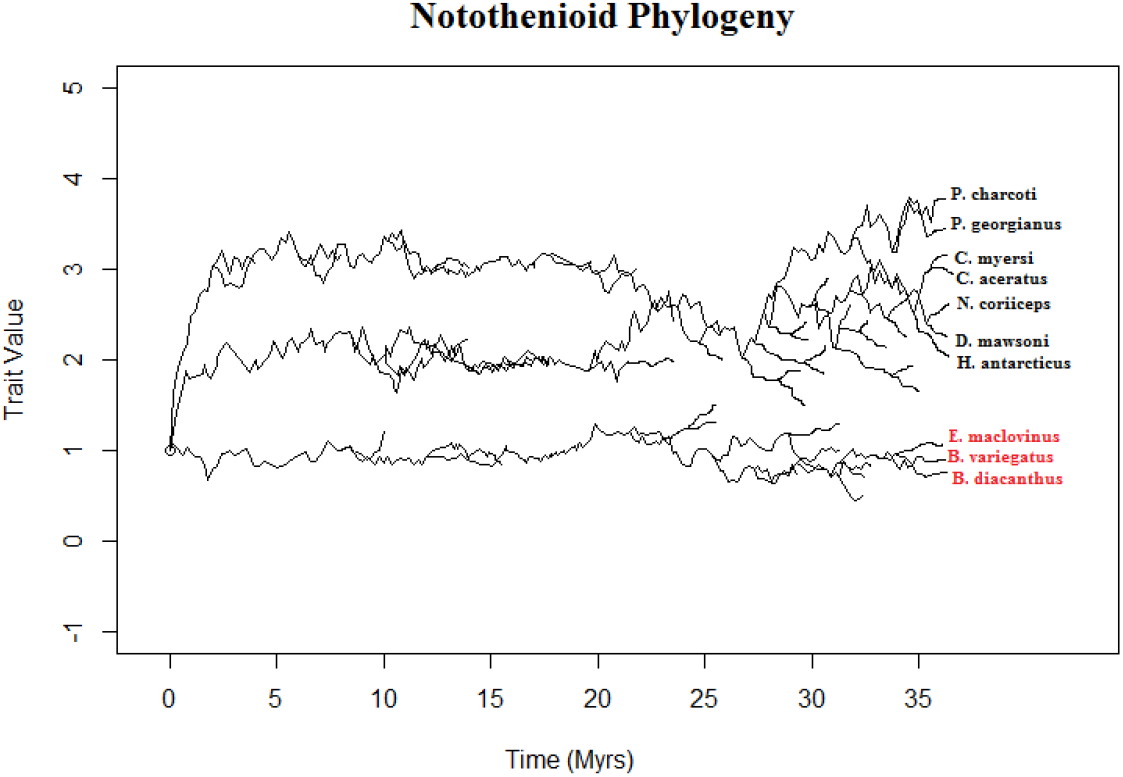
Simulated phylogeny of Antarctic notothenioids for last 35 million years, showing adaptive radiation during period of cooling of the Southern Ocean.

From the simulated phylogeny, we can also obtain a plot of the effective population size of notothenioids over time. The results are shown in Figure 6.

**Figure 6:**
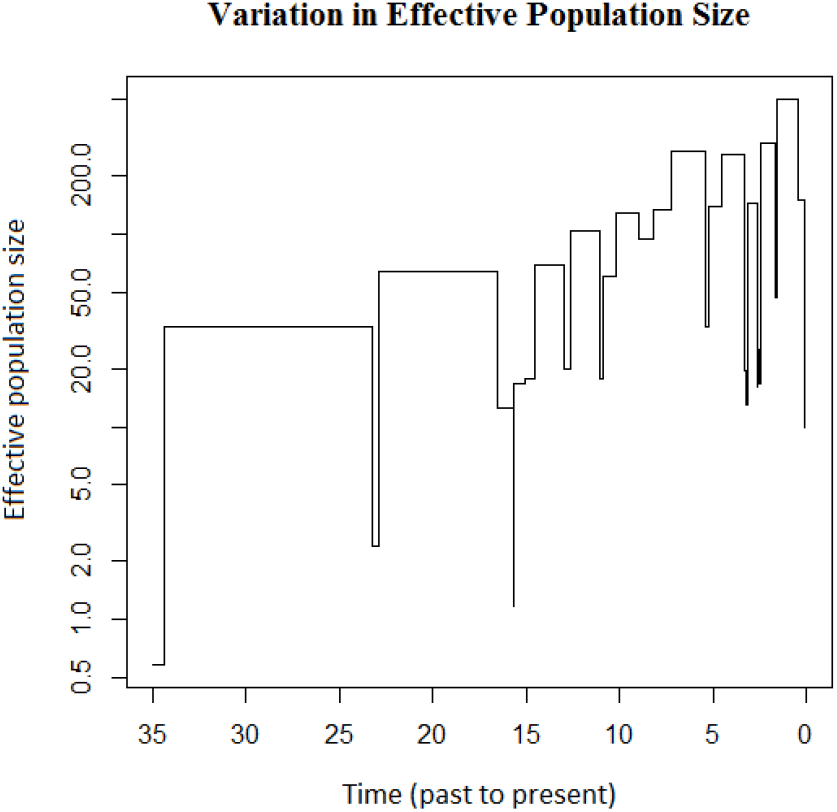
Variation in effective population size of Antarctic notothenioids over last 35 million years.

The effective population size (y-axis) is given as a relative measure, not in number of individuals. Time (x-axis) is given in millions of years ago. Note the population bottlenecks which appear in the last 5 million years, as a result of increased glacial cycles during this time.

## 4 Multivariate Simulation

The PhylSim package can also handle simulations with multiple trait values evolving together in time. In this case, the numerical trait *T* (*t*) would become 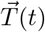, that is, a vector of trait values at time *t*, and the stochastic differential equation for the Ornstein-Uhlenbeck process becomes

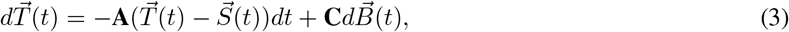

with **A** and **C** now as matrices acting on the corresponding vectors [2].

The solution for the multivariate case is therefore

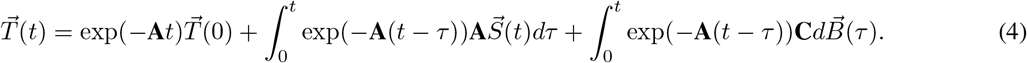

Hence, the above simulation could be run considering multiple kinase genes, or even entire gene families, to test for other hypotheses of evolution and adaptation. It is enough to define ‘X0’ to be a vector of trait values of interest, and to specify the evolutionary model to be a multivariate Ornstein-Uhlenbeck process, rather than single variable.

As an example, we consider the Antarctic species from the previous section, and simulate their diversification in the last 10 million years. The trait values we consider in the Ornstein-Uhlenbeck process are the number of copies of three kinase genes: Prkg1-201, Prkd3-201, and Mast3b-201. All three are important for intracellular signalling and response to oxidative stress, and have been extensively duplicated in Antarctic notothenioids, perhaps as an adaptation to the extreme environmental conditions in the Southern Ocean [7].

The results of the simulation are shown in Figure 7. We have included two additional species, *Trematomus bernacchii* (emerald notothen) [24], and *Pagothenia borchgrevinki* (bald notothen) [25], for which genetic data was also available.

**Figure 7:**
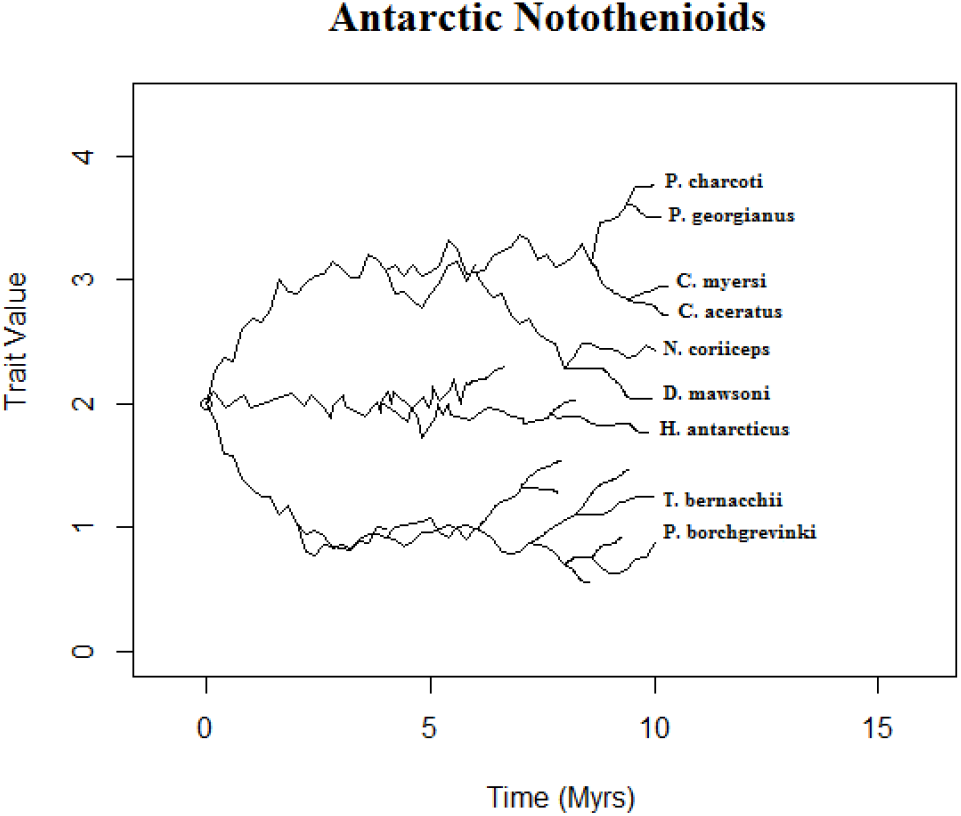
Simulated phylogeny of Antarctic notothenioids for last 10 million years using multiple kinase genes.

## 5 Parameter Estimation

We now consider parameter estimation using PhylSim. Let *θ* be the vector of true parameters of an evolutionary process on a phylogeny, and let *θ*′ be the estimated parameters. For results to be meaningful, we would like *E*[*θ*′] − *θ* to be zero, or at least below a certain tolerance.

We can calculate the probability that an estimator is unbiased using hypothesis testing [26]. In particular, we can take the null hypothesis to be

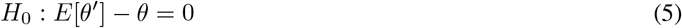

and

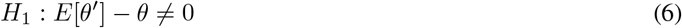

We use a one-way t-test with test statistic

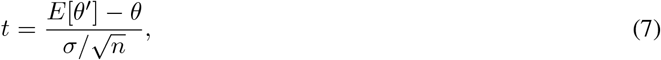

where *σ* is the sample standard deviation. Thus, if the p-value is below a value *α*, we reject the null hypothesis that the estimator is unbiased with significance level *α* [26]. In addition to the p-value, we can also consider the mean squared error (MSE) of the estimates, *E*[*θ*′ − *θ*]^2^, as another metric of estimator performance.

We start by simulating an evolutionary process with a vector of parameters *θ*_0_. We then use ABC inference to obtain the estimates of those parameters, 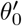 . Here, a variety of distance functions can be used in the ABC algorithm.

For distance between trait values, PhylSim uses the function ‘covmeandist’, which first estimates covariance matrices and mean vectors for the original and simulated data, and then computes the distance between them. It also has the function ‘covdist’, which only uses the covariance matrices to calculate the distance. For distance between trees, PhylSim uses functions provided by the ‘pcmabc’ package, namely ‘bdcoeffs’, ‘node heights’, and ‘logweighted node heights’ [3].

Different combinations of distance functions will give different p-values when testing a given evolutionary model. Hence, we can try different combinations to see which one performs best for each type of model.

Consider first a simple case, a univariate Brownian motion where *D* is the diffusion parameter. We simulate this process with a fixed value of *D* (in particular, *D* = 1), and then use the ABC algorithm with different distance functions to estimate this parameter. For each trial, we compute the p-value and mean squared error (MSE). After repeated simulation and estimation, we obtain the statistics shown in Figure 8.

**Figure 8:**
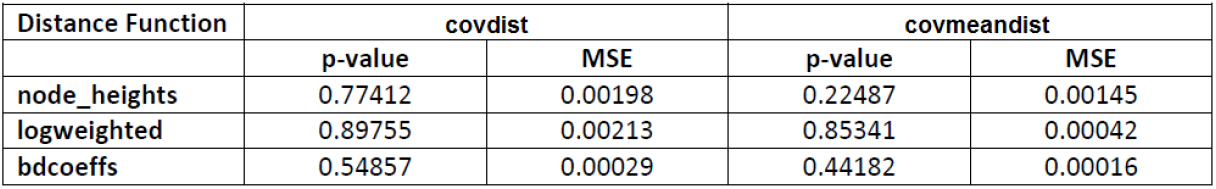
Summary of p-value and mean squared error (MSE) for different distance functions when estimating diffusion parameter *D* in a univariate Brownian motion.

From the table, we can see that the best combination of distance functions is ‘covmeandist’ for the distance between trait values, and ‘bdcoeffs’ for the distance between trees, as this combination has the smallest mean squared error for the estimate, while still having a sufficiently high p-value.

We apply the same approach to other evolutionary models to find the best combination in each case. The results are summarised in Figure 9. Note, however, that other combinations may also work well in certain scenarios, and that a more detailed study is necessary in order to draw any definitive conclusions.

**Figure 9:**
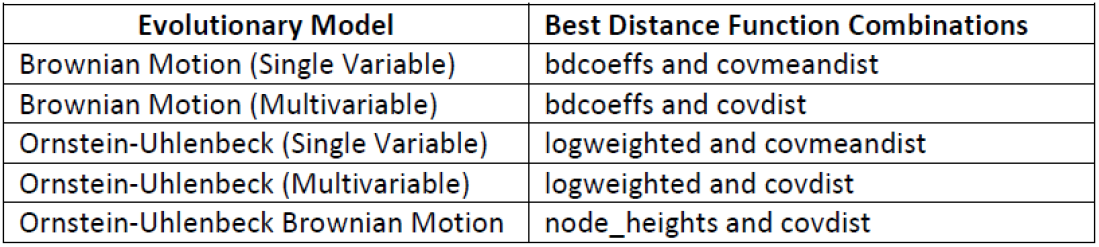
Summary of best distance function combinations for different evolutionary models.

## 6 Conclusions and Directions for Future Work

By allowing the user to vary speciation rates and evolutionary models over the course of a simulation, the PhylSim package provides a flexible tool for modeling trait evolution and speciation on a phylogeny. As a result, it can be used to test hypotheses of evolution and adaptation in scenarios where the speciation rate is believed to have varied significantly over evolutionary history.

We have considered the example of adaptive radiation of notothenioids in the Southern Ocean under changing climate conditions. As mentioned above, the simulation results produced by PhylSim agree with the findings of Near et al. (2012) [8], which suggest that the main diversification of Antarctic species occurred only in the last 5-10 million years, and not with the appearance of antifreeze proteins, which occurred much earlier [15].

Despite the robustness and flexibility of the PhylSim package, future work could be directed at increasing the range of evolutionary models that PhylSim can accept, beyond those provided by the ‘yuima’ package [4]. In addition, more complex speciation functions may be desirable for some applications, and this would also require some modification of the current implementation of PhylSim.

## 7 Acknowledgements

The authors wish to thank L.S. Peck (British Antarctic Survey) for providing valuable references and insight into the environmental history of Antarctica. We also wish to thank K. Bartoszek (Linkoping University) for his helpful comments and suggestions regarding the manuscript.

## References

[1] Butler, M.A., King, A.A. (2004). Phylogenetic comparative analysis: a modelling approach for adaptive evolution. Am. Nat. 164, 683695.

[2] Bartoszek et al. (2012). A phylogenetic comparative method for studying multivariate adaptation. Journal of Theoretical Biology 314, 2012 204215.

[3] Bartoszek, K., Lio, P. (2019). Modelling trait dependent speciation using Approximate Bayesian Computation. Acta Physica Polonica B Proceedings Supplement, 12(1), 2547.

[4] Brouste, A. et al. (2014). The YUIMA Project: A Computational Framework for Simulation and Inference of Stochastic Differential Equations. Journal of Statistical Software, 57(4), 151.

[5] Crame, J.A. (2018) Key stages in the evolution of the Antarctic marine fauna. Journal of Biogeography 45: 986994.

[6] Arvestad, L. et al. (2009). The gene evolution model and computing its associated probabilities. J. ACM, 56, 144.

[7] Clark et al. (2004). Antarctic genomics. Comparative and Functional Genomics 2004; 5: 230238.

[8] Near, T. J. et al. (2012). Ancient climate change, antifreeze, and the evolutionary diversification of Antarctic fishes. Proceedings of the National Academy of Sciences of the United States of America, 109, 34343439.

[9] Portner et al. (1998). Energetic aspects of cold adaptation: critical temperatures in metabolic, ionic and acid base regulation? In: Portner and Playle, Cold ocean physiology, Cambridge University Press, Cambridge, pp. 88–120.

[10] Akerborg, O. et al. (2009). Simultaneous Bayesian gene tree reconstruction and reconciliation analysis. Proc. Natl. Acad. Sci. USA, 106, 57145719.

[11] Clark and Peck (2009). HSP70 heat shock proteins and environmental stress in Antarctic marine organisms, Marine Genomics 2 2009, 1118.

[12] Peck, L. (2002). Ecophysiology of Antarctic marine ectotherms: limits to life, Polar Biology 2002, 25; 31–40.

[13] Buckley and Somero (2009). cDNA microarray analysis, Polar Biol 2009, 32:403415

[14] Sjostrand et al. (2012). DLRS: gene tree evolution in light of a species tree, Bioinformatics, Vol. 28 no. 22 2012, pages 29942995.

[15] DeVries, A.L.,Wohlshlag, D.E. (1969). Freezing resistance in some Antarctic fishes, Science 163, 10731075.

[16] Ahn, D.H., et al. (2017). Draft genome of the Antarctic dragonfish, Parachaenichthys charcoti, GigaScience, Volume 6, Issue 8, August 2017.

[17] Chen et al. (2019). The genomic basis for colonizing the freezing Southern Ocean revealed by Antarctic toothfish and Patagonian robalo genomes, GigaScience, Volume 8, Issue 4, April 2019.

[18] Shin, S.C., et al. (2014). The genome sequence of the Antarctic bullhead notothen reveals evolutionary adaptations to a cold environment, Genome Biol 15, 468.

[19] Kim, B., et al. (2019). Antarctic blackfin icefish genome reveals adaptations to extreme environments, Nat Ecol Evol 3, 469478.

[20] Bargelloni, L., et al. (2019). Draft genome assembly and transcriptome data of the icefish Chionodraco myersi reveal the key role of mitochondria for a life without hemoglobin at subzero temperatures, Commun Biol 2, 443.

[21] NCBI Sequence Read Archive, Accession Number ERX3357116.

[22] NCBI Sequence Read Archive, Accession Number ERX3357120.

[23] Berthelot, C., et. al. (2019). Adaptation of Proteins to the Cold in Antarctic Fish: A Role for Methionine?, Genome Biology and Evolution, Volume 11, Issue 1, January 2019, Pages 220231.

[24] Huth, T.J., Place, S.P. (2016). Transcriptome wide analyses reveal a sustained cellular stress response in the gill tissue of Trematomus bernacchii after acclimation to multiple stressors, BMC Genomics 17, 127.

[25] Bilyk, K., Cheng, C.H. (2014). RNA-seq analyses of cellular responses to elevated body temperature in the high Antarctic cryopelagic nototheniid fish Pagothenia borchgrevinki, Marine Genomics, Volume 18, Part B, December 2014, Pages 163–171.

[26] Wu, J. (2020). Comparing ABC distance functions for estimating parameters for phylogenetic comparative methods, Masters Project, IDA, Linkoping University, March 2020.

[27] Kutsukake, N., et al. (2014). Detecting phenotypic selection by approximate bayesian computation in phylogenetic comparative methods. In: -Modern Phylogenetic Comparative Methods and Their Application in Evolutionary Biology. Springer, pp. 409424.

